# Loss of GdpP function in *Staphylococcus aureus* leads to β-lactam tolerance and enhanced evolution of β-lactam resistance

**DOI:** 10.1101/2021.07.02.449101

**Authors:** Raymond Poon, Li Basuino, Nidhi Satishkumar, Aditi Chatterjee, Nagaraja Mukkayyan, Emma Buggeln, Liusheng Huang, Vinod Nair, Maria A. Argudín, Sandip K. Datta, Henry F. Chambers, Som S. Chatterjee

## Abstract

**Background:** We previously reported the presence of mutations in *gdpP* among *Staphylococcus aureus* strains that were obtained by serial passaging in β-lactam drugs. *gdpP* codes for a phosphodiesterase that cleaves cyclic-di-AMP (CDA), a newly discovered second messenger.

**Objectives:** We sought to identify the role of *gdpP* in β-lactam resistance of *S. aureus*.

**Methods:** CDA concentrations in bacterial cytosol were measured through mass-spectrometric analysis. *gdpP* deletion mutagenesis and their complemented strains were created in clinically relevant *S. aureus* strains to characterize its function.

**Results:** *gdpP* associated mutations among passaged strains were identified to cause loss of phosphodiesterase function, leading to increased CDA accumulation in the bacterial cytosol. Deletion of *gdpP* led to an enhanced ability of the bacteria to withstand a β-lactam challenge (two to three log increase in bacterial colony forming units) by promoting tolerance without enhancing MICs of β-lactam antibiotics. Our results demonstrate that increased drug tolerance due to loss of GdpP function can provide a selective advantage in acquisition of high-level β-lactam resistance and could lead to β-lactam treatment failure of *S. aureus* infections.

**Conclusions:** Loss of GdpP function increases tolerance to β-lactams that can lead to its therapy failure and can permit β-lactam resistance to occur more readily.

## Introduction

*Staphylococcus aureus* is an important pathogen that causes widespread nosocomial and community associated infections in humans.^1, 2^ Along with possessing a vast array of virulence factors, *S. aureus* also harbors the ability to display antibiotic resistance and antibiotic tolerance that enable bacteria to evade the action of antibiotics used for treatment.^3^ While resistance is typically characterized by an increase in the Minimum Inhibitory Concentration (MIC) for a target antibiotic, tolerance is a phenomenon where the survival of cells in presence of antibiotics is prolonged, without causing a change in drug MIC.^4^ β-lactams are an important class of antibiotics, which are often the first choice of treatment for *S. aureus* infections, primarily due to their high safety and efficacy. Over the past several decades, β-lactam resistance among contemporary clinical strains of *S. aureus* has called for the use of second-line agents that are often less safe and effective.^5^

β-lactam resistance in *S. aureus* is classically mediated by Penicillin Binding Protein 2a (PBP2a) and β-lactamase, which are encoded by *mecA* and *blaZ* respectively.^6^ While they have been well investigated, there is increasing evidence of the prevalence of non-classical mechanisms of β-lactam resistance.^7–9^ Our previous studies that were targeted towards identifying non-classical mechanisms of resistance, detected missense and nonsense mutations in *gdpP* among every resistant passaged strain.^10^ Recent clinical surveillance studies also detected similar *gdpP* associated mutations among several β-lactam non-susceptible isolates, reiterating their clinical relevance.^7, 11^ GdpP is a phosphodiesterase (PDE) that cleaves cyclic-di-AMP (CDA).^1, 12–15^ CDA is a newly discovered secondary messenger in bacteria and archaea that is synthesized from two molecules of ATP by the action of diadenylate cyclase (encoded by *dacA*).^16^ CDA has been deemed essential for bacterial cells and is implicated in a variety of cellular processes such as ion homeostasis, cell size and cell wall stress response among others in *S. aureus*.^17–20^ GdpP through its phosphodiesterase function helps maintain the intracellular levels of CDA through its cleavage. In this study, we focused on determining the role of *gdpP* associated mutations in surviving the antibacterial action of β-lactam drugs. Our results suggest that *gdpP* associated mutations detected among laboratory passaged and clinically isolated β-lactam resistant or non-susceptible strains of *S. aureus* were loss of function mutations that led to enhanced CDA concentrations in bacterial cytosol. Enhanced CDA concentrations facilitated bacterial survival against β-lactam antibiotics by promoting drug tolerance but did not alter drug MICs. β-lactam drug tolerance achieved through loss of GdpP function in *S. aureus* can lead to its therapy failure. Our findings further indicate that GdpP loss of function mutations arise very early during the evolution of β-lactam resistance and provide new insights into how attaining drug tolerance can promote faster evolution of β-lactam resistance in *S. aureus*.

## Materials and Methods

### Bacterial strains and growth conditions

Bacterial strains were grown in trypticase soy broth (TSB) as previously described.^14^ Allelic replacements and complementation were carried out with the plasmids *pKOR1* and *pTX_Δ_* respectively.^10^ Tetracycline resistant SF8300 (SF8300^tet^) was created by chromosomal integration of the plasmid *pLL29* as previously described.^21^ Bacterial strains, plasmids, and primers used in this study are listed in **Tables S1a, S1b, S2 and S3**, respectively.

### Antibiotic resistance measurements

MICs were determined by microbroth dilution method as described before.^22^ Briefly, 1×10^5^ CFU were incubated for 48 h at 37°C in 0.2 ml cation-adjusted Mueller Hinton broth (CAMHB) (BD Biosciences) containing increasing concentrations of antibiotic. MIC was recorded as the lowest concentration without growth at 48h.

Population analyses were done by the agar method as previously described.^14^ Tetracycline (12.5 mg/L) was added to the agar plates to select for the plasmid *pTX*_Δ_ whenever stated. Briefly, a 10 μl volume of serially diluted culture was spotted onto agar plates containing various concentrations of antibiotic. The plates were incubated at 37°C for 72 h. Plates were read and expressed as CFU ml^-1^. Ceftobiprole and ceftaroline were provided by Johnson and Johnson Pharmaceutical Research and Development and Forest labs respectively. All other antibiotics were purchased from Sigma-Aldrich.

### CDA measurement in bacterial cell lysates

CDA measurement was carried out as previously described ^23^ with a few modifications. Briefly, bacterial cells were collected after 6 h post culture, washed and lysed in PBS containing 1 mM EDTA using a FastPrep-24 homogenizer (MP Biomedicals). Bacterial cytosolic fraction was collected upon centrifugation. Aliquots of 40 μl cytosolic samples was mixed with 10 μl internal standard (20 ng/ml tenofovir), and 10 μl of the resulting mixture was injected into Liquid Chromatography-Tandem Mass Spectrometer (LC-MS/MS) system. The LC-MS/MS system consisted of AB Sciex API5000 tandem mass spectrometer, Shimadzu Prominence 20AD^XR^ UFLC pumps and SIL-20AC^XR^ autosampler. A hypercarb analytical column (100 x2.1mm, 3mm, thermos Sci. USA) and mobile phases 100 mM ammonium acetate (Buffer A) and acetonitrile (Buffer B) were used for separation. Electrospray ionization in negative ion mode as the ion source and multiple reactions monitoring with ion pairs m/z 657/133 for CDA and m/z 286/133 for the internal standard were used for quantification. Calibration standards were prepared with synthetic CDA (BioLog) dissolved in the PBS containing 1 mM EDTA. The calibration range was 10-500 ng/ml with lower limit of quantification at 10 ng/ml.

### Identification of GdpP function among clinical strains

We cloned the *gdpP* gene from the clinical strains that contained missense mutations with a constitutively expressing plasmid, *pTX_Δ_* and transformed the construct into a heterologous host, SF8300ex. The intracellular CDA abundance for each of these strains was measured through LC-MS/MS.

All other methods used in this study is presented in details in the supplementary text section.

## Results

### Mutations associated with *gdpP* cause loss of phosphodiesterase (PDE) activity in laboratory generated β-lactam resistant passaged strains and clinical isolates

Every resistant passaged strain previously obtained through passaging of COLnex and SF8300ex (lacking *mecA* and *blaZ*, the classical mediators of β-lactam resistance in wild-type COLn and SF8300 strains respectively) had a missense or nonsense mutation in *gdpP* that led to an amino acid alteration or a premature truncation of GdpP respectively **(Table S1a and Figure 1a)**.^1, 10, 12, 13, 15^ The nonsense mutations were such that they ablated the DHH-DHHA1 domains of GdpP, which mediate phosphodiesterase (PDE) activity to degrade CDA **(Figure 1a)**. Recent clinical surveillance studies have also identified similar *gdpP*-associated mutations among β-lactam non-susceptible *S. aureus* strains, further implicating their role in β-lactam resistance **(Table S4, Figure S1a)**.^7, 11^

**Figure 1:**
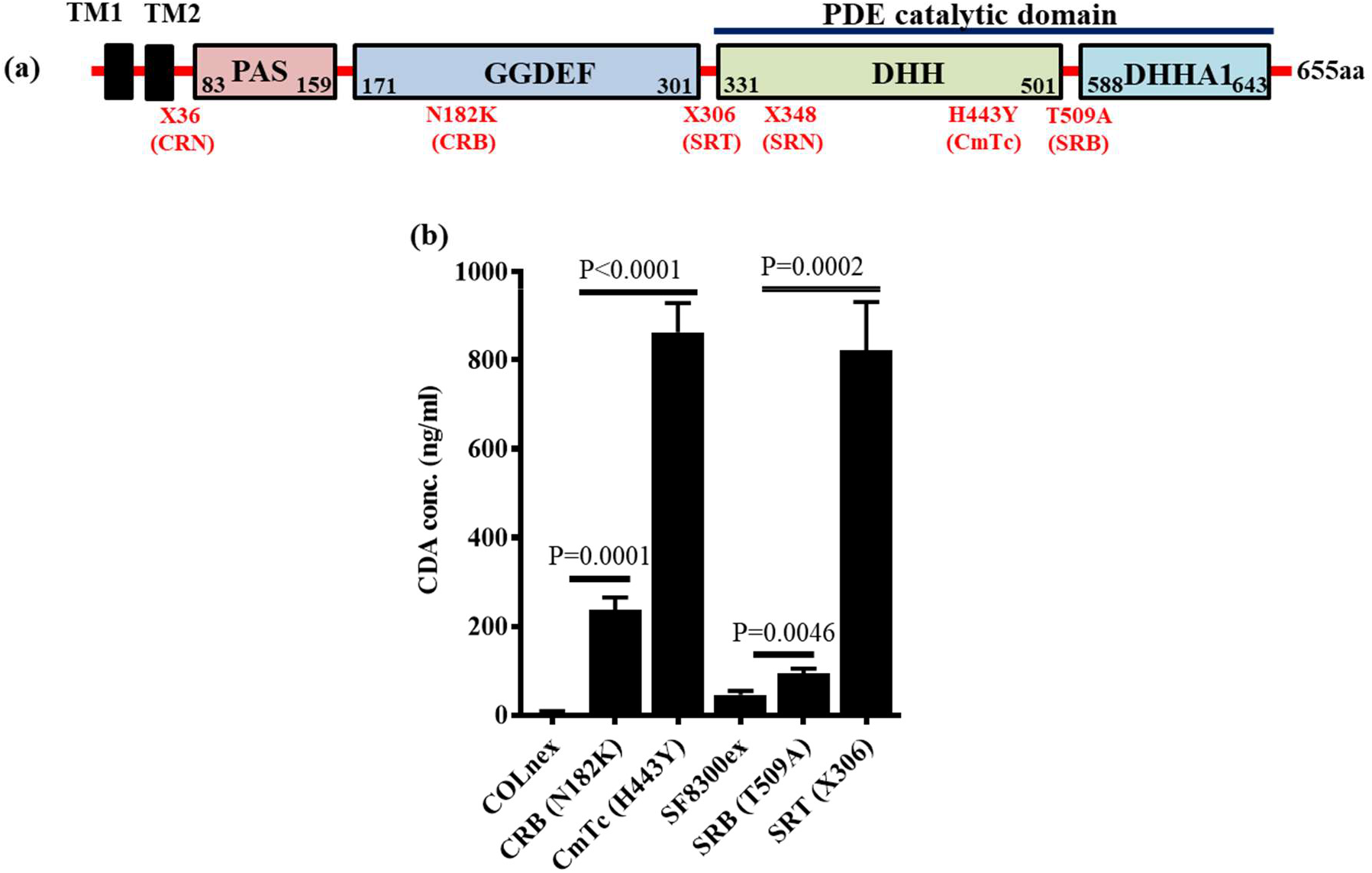
Passaged strains accumulated loss of GdpP function mutations. **(a)** Schematic diagram showing different domains of GdpP and mutations detected among passaged resistant strains. **(b)** Cyclic-di-AMP concentrations in the cytosol of passaged strains in comparison to their isogenic susceptible parental strains COLnex and SF8300ex. P values were obtained through two-tailed t-test analysis using GraphPad prism.

We hypothesized that the accumulated mutations of *gdpP* caused attenuated PDE function, **(Table S1a and Figure 1a)** leading to elevated CDA concentrations in passaged strains. To make this determination, we measured CDA abundance of the resistant passaged strains and compared them to their susceptible parents **(Table S1a)**. We evaluated a representative strain that had a nonsense mutation in *gdpP* (X306) that resulted in a premature truncation before the DHH-DHHA1 domains (SRT with X306; **Table S1a and Figure 1a**) and all the strains that accumulated missense mutations in our passaging study (CRB with N182K, CmTc with H443Y and SRB with T509A; **Figure 1a and Table S1a)**. We detected that every strain, including those having a missense mutation, displayed elevated levels of CDA compared to their isogenic parents, COLnex and SF8300ex **(Figure 1b)**. SRT had ~10-fold increased level of CDA. Notably, *gdpP* with the H443Y mutation (in the passaged strain CmTc) that substituted the 2^nd^ histidine residue of the DHH residues with a tyrosine also yielded similarly high levels of CDA in the bacterial cytosol. Two other *gdpP* mutants, CRB (N182K) and SRB (T509A) had significantly elevated CDA amounts compared to their isogenic wild-type strains **(Figure 1b)**. These results indicated that the accumulated mutations in *gdpP* altered GdpP’s phosphodiesterase activity that ranged from partial attenuation to complete loss of function. Similarly, the majority of *gdpP* mutations identified through a recent clinical surveillance study were GdpP loss of function mutations **(Table S4, Figure S1a)**. Of the 12 clinical strains analyzed for cytosolic CDA concentrations, eight were seen to have significantly increased CDA levels **(Figure S1b)**.

### Loss of GdpP function leads to a growth defect, enhanced ability to withstand β-lactam challenge but does not change MIC against β-lactam antibiotics

To determine the phenotypes associated with loss of GdpP function, *gdpP* was deleted from *S. aureus* SF8300, a clinical isolate that belongs to the USA300 background strain and from its isogenic *mecA* and *blaZ* excised variant, SF8300ex. This approach allowed us to evaluate the role of GdpP in isogenic background strains both with and without the classical mediators of β-lactam resistance. Deletion of *gdpP* caused increased accumulation of CDA in the bacterial cytosol and produced a growth defect compared to the isogenic wild-type strains **(Figure 2a, b)**. To assess if the observed Δ*gdpP*-related phenotypes were indeed due to loss of *gdpP*, we complemented the Δ*gdpP* mutants with constitutively expressing wild-type *gdpP*. We also transformed the wild-type and Δ*gdpP* mutant strains with an empty vector as control strains. *gdpP* complementation in Δ*gdpP* mutants restored their wild-type phenotypes **(Figure 2)**. Results of population analysis revealed that the Δ*gdpP* strains had at least a two to three log fold increase in the proportion of bacterial cells compared to their isogenic wild-type strains, indicating a heightened survival to β-lactam drug treatment **(Figure 2c)**. This also denoted that the observed phenotypes were attributed to *gdpP*. Since both *mecA* positive and negative strains (i.e. SF8300 and SF830ex respectively) displayed a similar pattern of β-lactam resilience, it indicated *mecA* (the major mediator of β-lactam resistance in *S. aureus*) likely did not play any important role in mediating the phenotypes that the Δ*gdpP* strains displayed **(Figure S2)**. In addition to the SF8300 Δ*gdpP* strain, we included another clinically relevant and epidemic-associated strain of *S. aureus*, MW2 (a member of the USA400 background) to confirm these findings **(Figure S3)**. These results reiterated the notion that GdpP played an important role in surviving β-lactam challenge in *S. aureus*.

**Figure 2:**
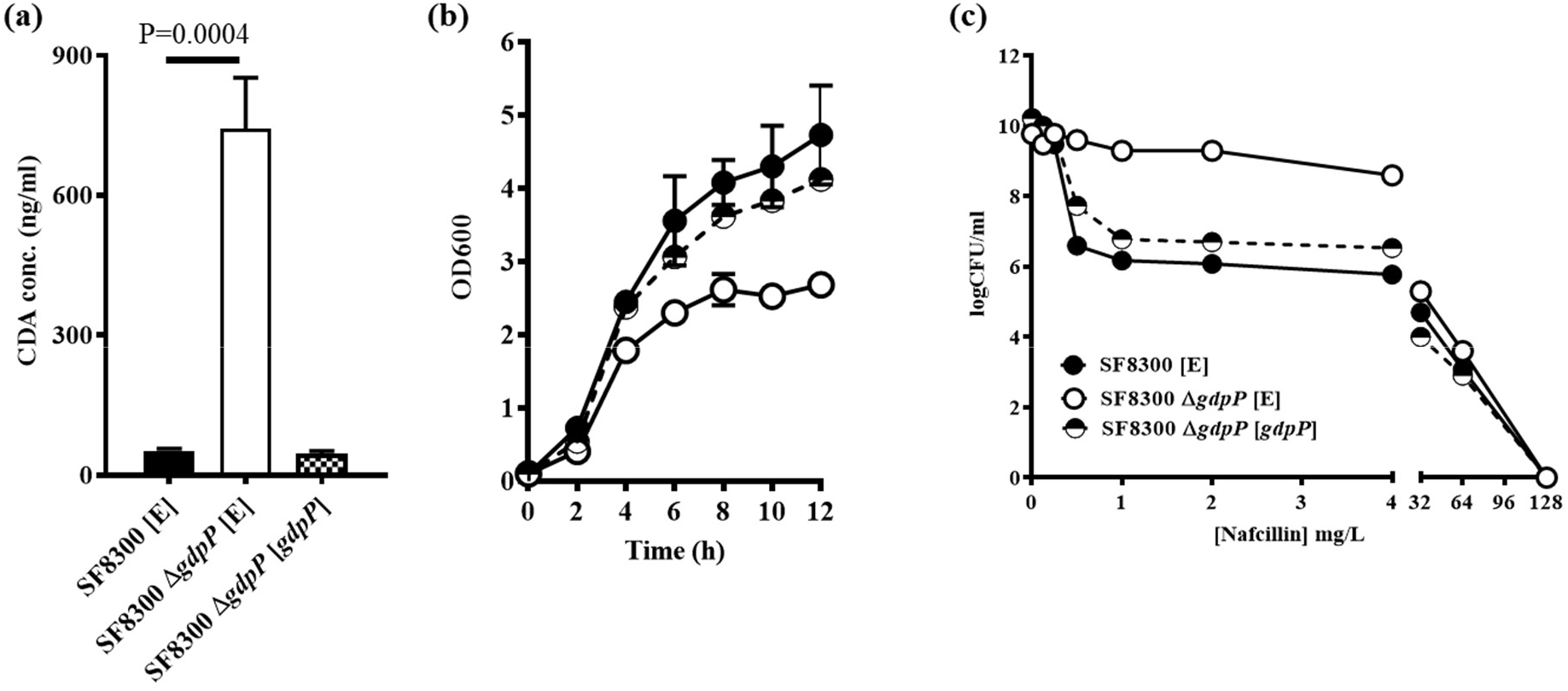
Complementation with *gdpP* restored wild-type phenotypes in SF8300 strains. **(a)** CDA levels in the cytosol **(b)** Growth pattern in TSB media **(c)** Population analysis with nafcillin of the complemented SF8300 and Δ*gdpP* strains. P values were obtained through two-tailed t-test analysis using GraphPad prism.

To further our understanding of the role of GdpP, we focused our study on *mecA* positive strains, which cause the majority of complicated and hard to treat *S. aureus* infections.^24^ To determine if CDA abundance alone was the factor driving the observed Δ*gdpP* associated β-lactam resilient phenotypes, the Δ*gdpP* mutant of SF8300 was complemented with *gdpP* mutants obtained from the passaged strains, namely CRB, CmTc, SRB and SRT **(Table S1a)**. The resultant isogenic strains displayed varying amounts of CDA **(Figure S4a)**. Growth assays **(Figure S4b)** and population assays **(Figure S4c)** showed a high correlation between CDA concentrations, bacterial growth attenuation and increased proportion of resistant populations.

Since the results of the population analysis showed a very similar endpoint of killing of the isogenic strains **(Figure S4c)**, we sought to determine if *gdpP*’s loss of function mediated enhanced β-lactam survival actually involved an MIC change. Results of the MIC assays did not show a substantial difference in MIC values between the strains **(Table 1)**. Thus, deletion of *gdpP* did not produce a resistant phenotype when measured through an MIC assay.

**Table 1:** Minimum Inhibitory concentrations (mg/L)

| Strains | NAF | CTX | FOX | CFZ | CPT |
| --- | --- | --- | --- | --- | --- |
| SF8300 | 32 | > 256 | 128 | 128 | 1 |
| SF8300 $\Delta gdpP$ | 32 | > 256 | 128 | 128 | 1 |

NAF- nafcillin; CTX- ceftriaxone; FOX- cefoxitin; CFZ- cefazoline; CPT- ceftaroline

### Loss of *gdpP* function leads to increased tolerance to β-lactams

Since deletion of *gdpP* allowed increased bacterial survival without a significant increase in the MIC values in presence of β-lactams, we determined if the *ΔgdpP* strains mediated β-lactam tolerance. We performed tolerance assays using two representative β-lactam antibiotics (nafcillin and cefoxitin). The Δ*gdpP* strains in SF8300 **(Figure 3)** and MW2 **(Figure S5)** backgrounds showed a classical^25^ tolerance response, displaying an increased survival over time compared to its isogenic wild-type strain following treatment of antibiotics. Complementation with *gdpP* in the Δ*gdpP* strains restored their wild-type phenotype. These results led to the suggestion that Δ*gdpP* strains were able to survive a β-lactam challenge via drug tolerance.

**Figure 3:**
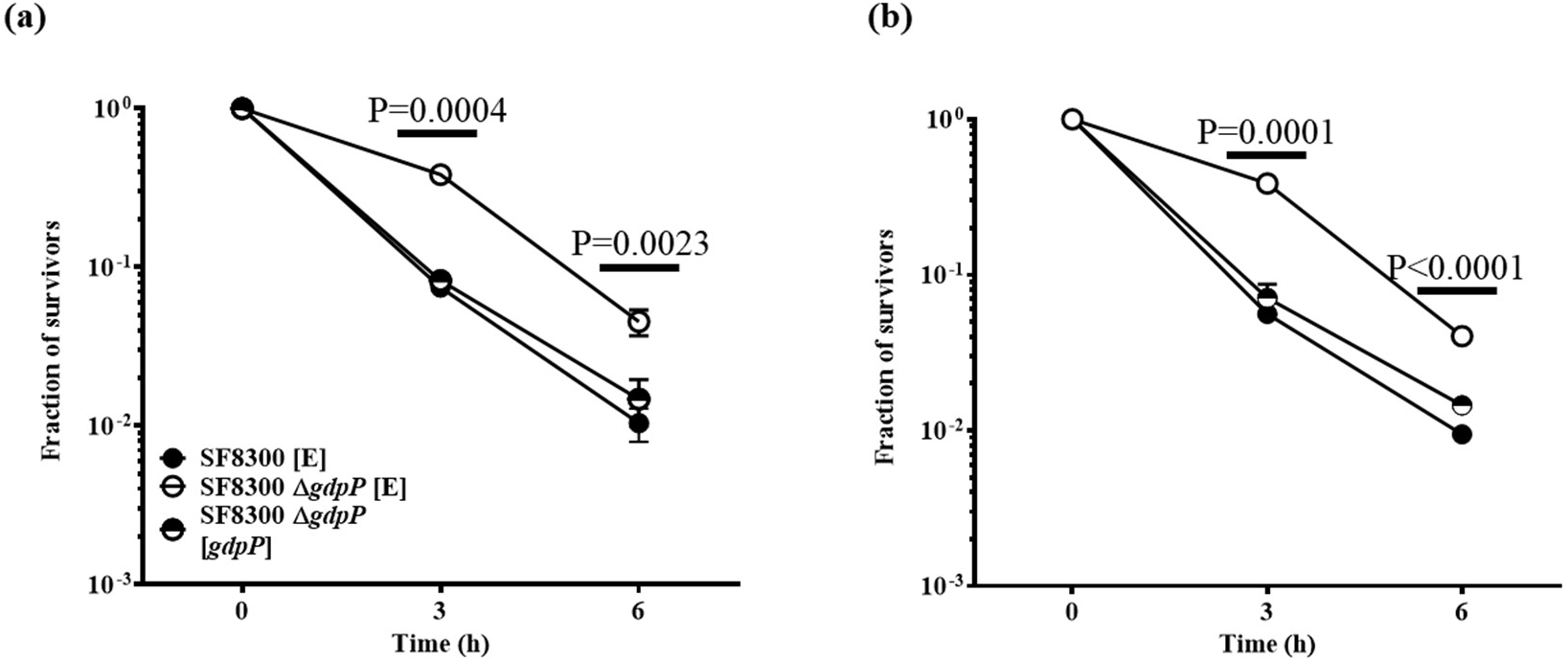
Deletion of *gdpP* leads to β-lactam tolerance in SF8300 strains. β-lactam tolerance assay carried out with **(a)** nafcillin (128 mg/L) and **(b)** cefoxitin (256 mg/L) P values represent difference between SF8300 [E] and SF8300 [*gdpP*] strains and were obtained through two-tailed t-test analysis using GraphPad prism.

### Loss of *gdpP* function mutations accumulate early in the process of β-lactam resistance and leads to faster evolution of β-lactam resistance

As mentioned above, the reported mutations in *gdpP* among resistant strains were obtained through serial passaging of the susceptible parental strains in presence of drugs that took several weeks to accomplish.^10^ While whole-genome sequencing enabled the identification of mutated genes, our study design did not inform about the specific stage of the passage at which the mutations were acquired. To determine this, we sequenced *pbp4* and *gdpP* in one of the passaged strain SRT,^14^ for which bacterial stock cultures from each day of passaging were available. SRT was obtained by passaging the susceptible SF8300ex (*mecA* and *blaZ* excised) strain in increasing concentrations of ceftaroline for 22 days. At day 22, SRT was able to grow in bacterial media containing 256 μg/ml ceftaroline.^14^ The purified SRT clone that was used for subsequent studies showed high-level resistance to β-lactam drugs and had *pbp4* promoter and gene mutations in addition to having the X306 mutation in GdpP **(Figure 1a)**.^10^ Bacteria from the daily stock cultures of SRT passage were streaked onto TSA plates and three colonies from each day of passage were randomly chosen for sequencing of *gdpP* and *pbp4* promoter and gene. We detected that SRT acquired its *gdpP*-associated X306 mutation as early as on day 4 of passaging during which bacteria were grown in media containing 0.5 μg/ml of ceftaroline. On day 4, SRT did not have any *pbp4* related mutations suggesting that *gdpP* associated mutations were acquired very early during the evolution of resistance in SRT. Of note, we also detected *gdpP* mutations among passaged strains obtained in day 3, which were different from the X306 detected on day 4 **(Table S5)**.

To determine if tolerance mediated by loss of GdpP function could facilitate evolution of β-lactam resistance, we performed passaging of the susceptible SF8300ex and its isogenic Δ*gdpP* strains in two β-lactam antibiotics (nafcillin and cefoxitin). Results of this study revealed that the Δ*gdpP* strain was able to develop resistance faster compared to its isogenic SF8300ex strain **(Figure S6)**. Thus, GdpP-mediated tolerance can lead to faster evolution of resistance.

### Loss of *gdpP* function could lead to β-lactam treatment failure

Since the therapeutic outcome of β-lactam tolerance due to loss of GdpP function is unknown, we created a tetracycline resistant SF8300 strain (SF8300^tet^) by integrating *pLL29* (single copy chromosomal integration vector) in its genome.^21^ The resultant SF8300^tet^ strain displayed no difference in β-lactam resistance phenotypes when measured through MIC assays **(Table S6)** or population analysis **(Figure S7a)**. There was also no change in CDA amounts in the bacterial cytosol **(Figure S7b)** when compared to its isogenic wild-type strain, SF8300. To test the impact of Δ*gdpP* induced tolerance on β-lactam treatment outcome, SF8300^tet^ was mixed with SF8300 Δ*gdpP* strain in a 1:1 ratio and was treated with different nafcillin concentrations. The resultant bacterial mixture was incubated for 24 h upon which surviving bacterial CFUs were enumerated by plating them onto TSA plates. The bacterial titers of this assay at 24 h showed a stepwise decrease of total surviving bacteria in a nafcillin concentration-dependent manner **(Figure 4a)**. To determine the respective abundance of bacterial strains in the mixture, the surviving bacteria were selected onto TSA and TSAtet (TSA containing tetracycline) plates. The results of this analysis showed a preponderance of the SF8300 Δ*gdpP* strain (i.e. tetracycline sensitive colonies) in the mixture **(Figure 4b)**. Furthermore, the Δ*gdpP* strain outcompeted the wild-type bacteria in a nafcillin concentration dependent manner as well. These results suggested that the Δ*gdpP* strain could survive a β-lactam challenge better than its isogenic wild-type strain. The results of these experiments also revealed that at 24 h following incubation, the fraction of Δ*gdpP* strain was slightly attenuated compared to that of the wild-type strain without any antibiotic treatment, probably due to the inherent growth defect that the Δ*gdpP* strain displayed. This suggested that tolerance mediated by *gdpP* could indeed lead to β-lactam non-susceptibility or therapy failure.

**Figure 4:**
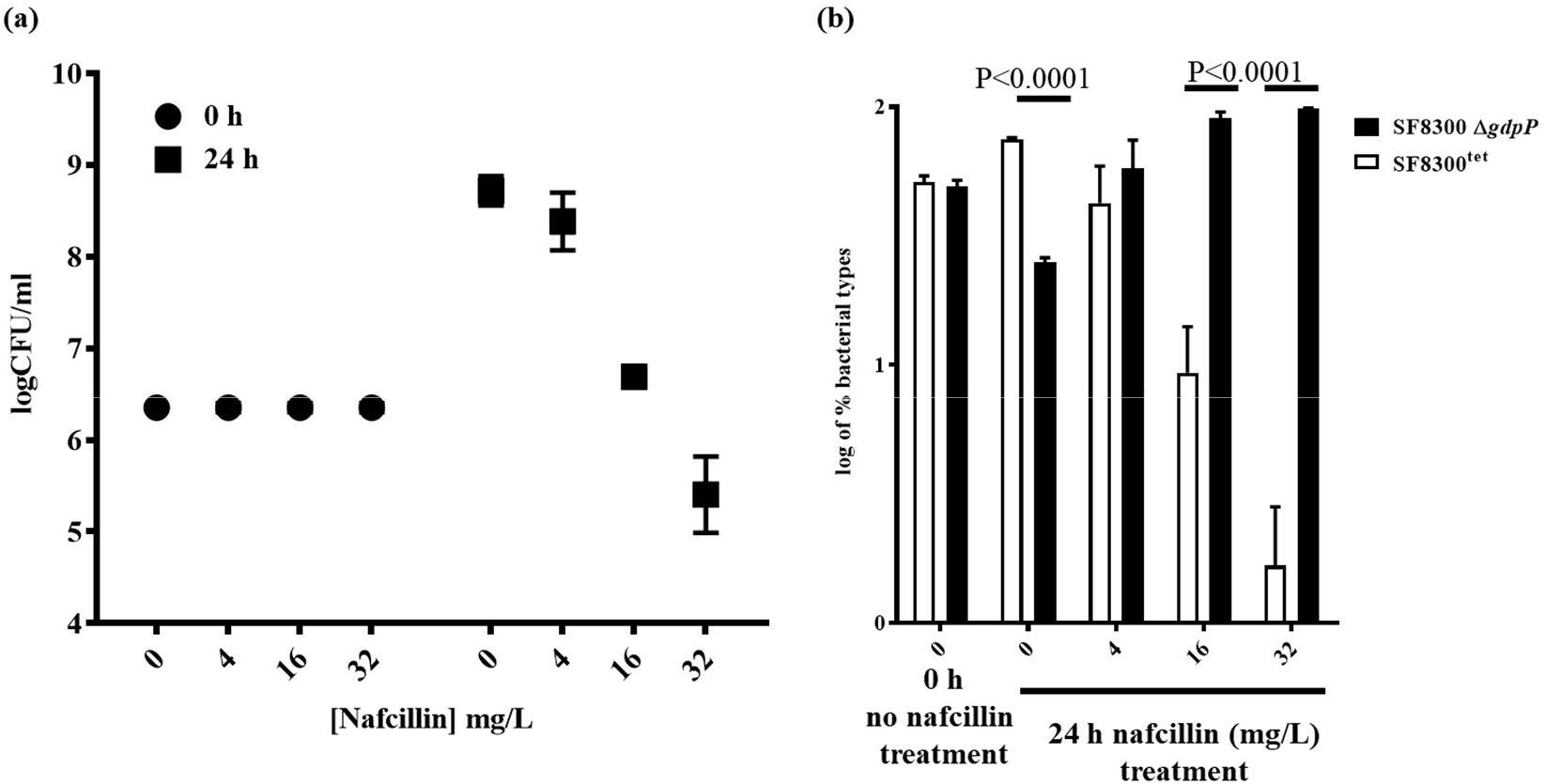
Deletion of *gdpP* leads to β-lactam resistance. **(a)** Survival of 1:1 mixed co-cultures of SF8300^tet^ and SF8300 Δ*gdpP* strains upon nafcillin challenge **(b)** Fraction of SF8300^tet^ and SF8300 Δ*gdpP* strains in the mixture

## Discussion

Previous work performed in Gram-positive firmicutes such as *Bacillus subtilis^26^, Listeria monocytogenes*^27^ and *S. aureus*^17, 28^ among others have shown a broad role of CDA in controlling bacterial physiology including osmoregulation, peptidoglycan metabolism, biofilm formation and virulence and have also drawn associations between CDA levels to β-lactam resistance of bacteria. While reduction of CDA concentration (through inhibition of its synthesis machinery) in bacterial cells has been shown to sensitize bacteria to β-lactams, increased amount of CDA (via inhibition of its degradation machinery) produces resistance to β-lactam drugs^17^. The mechanism/s through which CDA modulates drug sensitivity/resistance however remains currently unknown.

A high-level association in detection of mutations in *gdpP* (the machinery that degrades CDA) among laboratory generated and natural strains of *S. aureus* that were non-susceptible or resistant to β-lactam drugs prompted us to determine their role in surviving β-lactam drug challenge.^7, 29^ Our results showed that the *gdpP* mutations, by virtue of their loss of function enabled increased intracellular CDA concentration in *S. aureus* **(Figure 1b, Figure S1b)**, which in turn provided a significant fitness advantage in surviving β-lactam challenge in a manner that was dependent on CDA concentration. Experiments carried out to decipher an enhanced understanding of CDA’s role enabling better survival to a drug challenge indicated that increased CDA concentrations promoted β-lactam tolerance without altering drug resistance when measured through classical means such as MIC or population analysis **(Table 1, Figure 2c)**. Although population analysis results showed homo-resistance characteristics of bacterial strains with high CDA concentrations **(Figure 2c, Figure S3c)**, they indicated an almost identical endpoint of bacterial killing to that of wild-type bacteria. Thus, the term “resistance” in previous studies that associated CDA levels with drug resistance might have been used loosely. Antibiotic tolerance has recently emerged as an important factor that is distinct from traditional resistance, which reduces drug efficacy by producing slower killing of bacteria upon chemotherapeutic challenge.^4, 25, 30^ Indeed our results show that acquisition of β-lactam tolerance through GdpP’s loss of function could lead to β-lactam therapy failure due to slower killing and thereby complicating treatment outcome **(Figures 3 and 4)**.

Recent studies have reported that increased *mecA* expression can enable *S. aureus* strains to display a β-lactam homo-resistance phenotype,^28^ similar to that observed among strains with elevated CDA concentrations. Western blot analysis did not indicate altered PBP2a expression among isogenic Wt and Δ*gdpP* strains used in this study **(Figure S8a)**. The results of bocillin assay further showed unaltered expression of housekeeping PBPs (PBP 1 through 4) suggesting that they were also likely not involved in GdpP mediated β-lactam tolerance **(Figure S8b)**. We created two additional isogenic set of strains (COLn and N315) to verify these results since the MW2 Δ*gdpP* strain showed slight overexpression of PBP4 in the bocillin assay. Complementation with *gdpP* did not alter the PBP4 expression of MW2 Δ*gdpP* strain **(Figure S8c)** suggesting that the elevated PBP4 level is likely due to secondary site mutation/s in this strain. Appearance of similar homo-resistance phenotypes displayed by SF8300 and SF8300ex strains **(Figure S2)** along with detection of loss of GdpP function mutations among both MRSA and MSSA natural strains^7, 11^ **(Table S4, Figure S1b)** further supported the notion that GdpP mediated effects are most likely carried out through mechanisms that are independent of bacterial PBPs.

Appearance of GdpP loss of function mutation among one of the resistance-passaged strain, SRT, showed that its acquisition preceded mutations that enabled antibiotic resistance indicating that tolerance precedes resistance during evolution of antibiotic resistance **(Table S5)**. It also indicated that antibiotic tolerance is an independent biological phenomenon than that of resistance, which potentially employs separate mechanism/s to aid antibiotic survival^25^. This is likely the reason why GdpP’s loss of function mutations could be identified in natural strains that are non-susceptible but not resistant to β-lactam drugs.^7^

The mechanism through which GdpP’s loss of function enables β-lactam tolerance is unknown. Slower growth rate of the strains with elevated CDA levels does not seem to cause β-lactam tolerance as both diminished and elevated CDA levels, which oppositely affects drug tolerance, reportedly inhibit bacterial growth.^19^ Thus, slower growth rate likely precludes β-lactam tolerance. Apart from a slightly smaller cell size, results of electron microscopic studies did not show much difference in bacterial gross cell morphologies including that of cell wall among the isogenic Wt and *ΔgdpP* strain pairs used in our study **(Figure S9)**. This ruled out the possibility that an altered cell wall structure is the contributing factor for β-lactam tolerance, although further studies are required in order to determine this with more certainty. Previous studies have shown that CDA acts as a vital gatekeeper of ion transporting channels (most prominently that of potassium transporters, KdpFABC and KtrAB/AD systems)^18, 31^ and thereby controlling ion homeostasis in bacterial cells. It is thus imperative that elevated CDA concentrations due to GdpP’s loss of function would lead to reduced potassium ion concentrations in bacterial cells, which in turn would increase bacterial turgor pressure forcing bacteria to alter their cell structure and cell wall stiffness. Indeed, similar to our observation, GdpP loss of function mutants have previously been shown to display 30% smaller staphylococcal cell size.^17^ Since changes in cell shape in bacteria have recently been shown to influence antibiotic efficacy,^32^ it will be tempting to determine if changes in physical parameters such as cell shape or stiffness brought in by CDA mediated ionic imbalance could contribute to β-lactam tolerance.

## Acknowledgments

We would like to thank Joseph Schaefer and Aubre Gilbert for their technical help. SSC would like to thank Dr. Nick Carbonetti for critically reading the manuscript. SSC supervised LB and EB while he was at UCSF.

## Funding statement

This work was funded by NIH grants 2R01AI100291 and R21AI142501 and startup funds provided by the University Systems of Maryland to SSC.

## Transparency declaration

The authors declare no conflict of interests.

## Notes

### Competing Interest Statement

The authors have declared no competing interest.

