## Supplementary material for "Loss of GdpP function in *Staphylococcus aureus* leads to β-lactam tolerance and enhanced evolution of β-lactam resistance"

**Supplementary text:**

**Materials and methods:**

Construction of *gdpP* deletion mutant and its complemented strains: To create *gdpP* deletion knockout strains, up- and down-stream regions of *gdpP* were PCR amplified with the primers *gdpP*-P1 & P2 and *gdpP*-P3 & P4, respectively. The full-length *gdpP* gene was PCR amplified using splice overlap extension PCR and was cloned into *pKOR1*. The resultant plasmid was sequence confirmed and transformed to the target strains. Allelic replacement of *gdpP* was carried out as previously described.[1] The deletion mutation was confirmed by PCR and through analytical sequencing.

To create the constitutively expressing *gdpP* in $pTX_{\Delta}$, full-length *gdpP* gene was PCR amplified from the genomic DNA of strains as mentioned in three different fragments using PCR primers *gdpP*-1 & 2, 3 & 4 and 5 & 6. The full-length *gdpP* was obtained by performing splice overlap extension PCR with the PCR fragments that were obtained in the previous step. The fragmented PCR amplification of the full-length *gdpP* was necessary to remove additional BamH1 & Mlu1 internal cutting sites, which were the restriction enzymes used to clone the full-length *gdpP* into the constitutively expressing vector $pTX_{\Delta}$. The resultant plasmid was sequence verified for sequence accuracy of the cloned *gdpP*s.

*gdpP*s from the selected clinical isolates **(Table S4)** were amplified and cloned into the $pTX_{\Delta}$ plasmid. The plasmids were first transformed into RN4220 followed by transduction into SF8300ex *ΔgdpP*.

Tolerance assay: Bacterial tolerance assay was carried out as previously described.[2] Briefly, 1x $10^6$ bacteria from overnight cultures were inoculated in 10 ml TSB with the indicated drug.

Bacteria were incubated at 37°C in a shaking incubator and plated at 0, 3 and 6 h for enumeration of CFUs.

Competition assay: Bacterial inoculum (SF8300$^{tet}$ and SF8300 Δ*gdpP* strains) was prepared by centrifugation of 10 ml bacterial culture that was grown in TSB media for 24 h followed by wash with PBS containing 10% glycerol. The bacterial pellet was re-suspended in PBS containing 10% glycerol, aliquoted in 1ml batches and stored at -80ºC. Bacterial titers were determined by plating serial dilutions of bacteria from 2 different tubes of stored bacteria. For competition assay, 2X $10^6$ bacteria (1:1 mixture of the 2 strains mentioned above) were inoculated in 10 ml TSB. The following conditions were tested 1) No drug and 2) nafcillin treatment (4, 16 and 32 mg/L). Bacterial CFU were determined by plating bacteria on TSA plates at 0 and 24 h post inoculation. Enumeration of respective number of SF8300$^{tet}$ and SF8300 Δ*gdpP* strains were determined by replica plating the bacterial colonies onto TSA and TSA containing tetracycline (3 mg/L).

Western Blotting: Overnight cultures of bacteria were subcultured into 50mL flasks containing TSB such that the initial $OD_{600}$ of the flasks were 0.1. The flasks were cultured for 2 h and 2 mg/L nafcillin was added. Bacterial cells were harvested following 2 h incubation of antibiotic addition. The collected cells were washed with Phosphate Buffered Saline (PBS), resuspended in lysis buffer (50mM Tris pH 7.5, 150mM NaCl, 2mM EDTA, 1mM Sodium Pyrophosphate, 1X CompleteMini protease inhibitor cocktail (Roche)) and were mechanically lysed using the FastPrep FP120 (Thermo Savant). Following centrifugation, the supernatant was collected and cell membrane fractions were isolated by performing ultracentrifugation (66000g, Sorvall WX Ultra 80 Centrifuge, Thermo Fisher Scientific). After resuspending the pellet with lysis buffer, protein estimation was carried out using Pierce BCA Protein Assay kit (Thermo Fisher). Western blot assay was carried out with Rabbit polyclonal anti-PBP2a antibody (RayBiotech) at a 1:1000 dilution for overnight incubation and Goat anti-Rabbit secondary antibody (Azure Biosystems) (1:10000) for 1 h.

Bocillin assay: As previously described,[3] overnight cultures of the selected *S. aureus* strains were diluted in TSB such that the $OD_{600}$ value was 0.1. Cells were grown at 37°C, 180 rpm until $OD_{600}$ of 1 after which 50 mL of the culture was collected for centrifugation at 2451 g (3500 rpm) to obtain cell pellets. The pellets were dried and resuspended in 500 μL PBS and were then mechanically lysed using the FastPrep FP120 (Thermo Savant). Following centrifugation, the supernatant was collected and cell membrane fractions were isolated by performing ultracentrifugation (66000g, Sorvall WX Ultra 80 Centrifuge, Thermo Fisher Scientific). The obtained pellets were resuspended in 100 μL PBS and protein estimation was carried out using Pierce BCA Protein Assay kit (Thermo Fisher). The membrane fraction was then incubated with 10 μM Bocillin-Fl (Thermo Fisher) for 30 minutes at 35°C. The reaction was stopped by boiling with sample buffer for 10 minutes following which 20 μg of protein was loaded onto a 10% SDS-polyacrylamide gel and electrophoresis was carried out at 80V. The bands were visualized using the Typhoon 9410 imager (Amersham/GE Healthcare).

Electron microscopy: Transmission Electron Microscopy (TEM) was carried out as described previously.[4] Briefly, overnight bacterial cultures were collected and fixed with 2.5% glutaraldehyde in 0.1M sodium cacodylate buffer (Electron Microscopy Sciences). Following post-fixation with 1% osmium tetroxide reduced with 0.8% potassium ferrocyanide, the cell pellets were treated with 1% tannic acid and stained with uranyl acetate replacement (UAR). Samples were dehydrated with a graded ethanol series, followed by dehydration with acetone and embedded in Embed 812 resin. Thin sections were cut with a Leica EM UC6 ultramicrotome (Leica), and stained with 1% UAR and Reynold's lead citrate prior to viewing at 120 kV on a Tecnai BT Spirit transmission electron microscope (FEI). Digital images were acquired with a Hamamatsu side mount digital camera system (Advanced Microscopy Techniques).

For scanning electron microscopy, cell suspensions were adsorbed on silicon chips and fixed with 2.5% glutaraldehyde in 0.1m sodium cacodylate buffer. Similar to the TEM specimen processing, the cells were post-fixed with 1% reduced osmium tetroxide and dehydrated in graded ethanol series. The specimen was dried using a critical point dryer (Bal-Tech AG), sputter coated with iridium (South Bay Technology, Inc.) and imaged on a Hitachi SU8000 scanning electron microscope (Hitachi High Technologies).

<u>Sequencing</u>: Fidelity of all the mutants and plasmid constructs were validated through Sanger sequencing (Eurofins Genomics, USA).

<u>Bioinformatics and statistical analysis</u>: Statistical analyses were performed using GraphPad Prism. Comparisons between groups were analyzed two-tailed t-test whenever stated. DNA sequence analysis was performed using DNAstar software.

**Supplementary tables and figures**

**Table S1a: List of wild-type and passaged strains used in this study**

| Strains | Notes | *pbp4* promoter | PBP4 | GdpP |
|---|---|---|---|---|
| COLnex | *mecA* excised COLn[5] | - | - | - |
| CRB | COLnex passaged in Ceftobiprole[5] | 36bp insertion | E183A; F241R | N182K |
| CmTc | COLnex passaged in Ceftaroline[6] | A>C at -399bp upstream of *pbp4* start codon | T201A; F241L | H443Y |
| SF8300ex | *mecA* and blaZ excised SF8300[6] | - | - | - |
| SRB | SF8300ex passaged in Ceftobiprole[6] | - | E183V; F241R | T509A |
| SRT | SF8300ex passaged in Ceftaroline[6] | "A" deletion at -378bp upstream and 11bp deletion at 300bp upstream of pbp4 start codon | N138K; H270L | X306 |
| SF8300 | Community associated MRSA | - | - | - |
| MW2 | Community associated MRSA | - | - | - |
| COLn | Hospital associated MRSA | - | - | - |
| N315 | Hospital associated MRSA | - | - | - |
| RN4220 | Laboratory *S. aureus* strain | - | - | - |

**Table S1b: List of strains created in this study**

| Strains created in this study | Notes |
|---|---|
| SF8300 Δ*gdpP* | *gdpP* deleted SF8300 strain |
| SF8300ex Δ*gdpP* | *gdpP* deleted SF8300ex strain |
| MW2 Δ*gdpP* | *gdpP* deleted MW2 strain |
| COLn Δ*gdpP* | *gdpP* deleted COLn strain |
| N315 Δ*gdpP* | *gdpP* deleted N315 strain |
| SF8300 [E] | SF8300 with empty constitutively expression vector $pTX_{\Delta}$ |
| SF8300 Δ*gdpP* [E] | SF8300 Δ*gdpP* with empty constitutively expression vector $pTX_{\Delta}$ |
| SF8300 Δ*gdpP* [*gdpP*] | SF8300 Δ*gdpP* with $pTX_{\Delta}$ expressing gdpP from SF8300 |
| MW2 [E] | MW2 with empty constitutively expression vector $pTX_{\Delta}$ |
| MW2 Δ*gdpP* [E] | MW2 Δ*gdpP* with empty constitutively expression vector $pTX_{\Delta}$ |
| MW2 Δ*gdpP* [*gdpP*] | MW2 Δ*gdpP* with $pTX_{\Delta}$ expressing *gdpP* from SF8300 |
| SF8300 Δ*gdpP* [CRB] | SF8300 Δ*gdpP* with $pTX_{\Delta}$ expressing *gdpP* from CRB |
| SF8300 Δ*gdpP* [CmTc] | SF8300 Δ*gdpP* with $pTX_{\Delta}$ expressing *gdpP* from CmTc |
| SF8300 Δ*gdpP* [SRB] | SF8300 Δ*gdpP* with $pTX_{\Delta}$ expressing *gdpP* from SRB |
| SF8300 Δ*gdpP* [SRT] | SF8300 Δ*gdpP* with $pTX_{\Delta}$ expressing *gdpP* from SRT |
| SF8300ex *ΔgdpP* [MSSA 4] | SF8300ex Δ*gdpP* with $pTX_{\Delta}$ expressing *gdpP* from MSSA 4 |
| SF8300ex *ΔgdpP* [MSSA 7] | SF8300ex Δ*gdpP* with $pTX_{\Delta}$ expressing *gdpP* from MSSA 7 |
| SF8300ex *ΔgdpP* [MSSA 9] | SF8300ex Δ*gdpP* with $pTX_{\Delta}$ expressing *gdpP* from MSSA 9 |
| SF8300ex *ΔgdpP* [MSSA 13] | SF8300ex Δ*gdpP* with $pTX_{\Delta}$ expressing *gdpP* from MSSA 13 |
| SF8300ex *ΔgdpP* [MSSA 19] | SF8300ex Δ*gdpP* with $pTX_{\Delta}$ expressing *gdpP* from MSSA 19 |
| SF8300ex *ΔgdpP* [MSSA 21] | SF8300ex Δ*gdpP* with $pTX_{\Delta}$ expressing *gdpP* from MSSA 21 |
| SF8300ex *ΔgdpP* [MSSA 22] | SF8300ex Δ*gdpP* with $pTX_{\Delta}$ expressing *gdpP* from MSSA 22 |
| SF8300ex *ΔgdpP* [MSSA 24] | SF8300ex Δ*gdpP* with $pTX_{\Delta}$ expressing *gdpP* from MSSA 24 |
| SF8300ex *ΔgdpP* [MSSA 25] | SF8300ex Δ*gdpP* with $pTX_{\Delta}$ expressing *gdpP* from MSSA 25 |
| SF8300ex *ΔgdpP* [MSSA 27] | SF8300ex Δ*gdpP* with $pTX_{\Delta}$ expressing *gdpP* from MSSA 27 |
| SF8300ex *ΔgdpP* [MSSA 28] | SF8300ex Δ*gdpP* with $pTX_{\Delta}$ expressing *gdpP* from MSSA 28 |
| SF8300ex *ΔgdpP* [MSSA 32] | SF8300ex Δ*gdpP* with $pTX_{\Delta}$ expressing *gdpP* from MSSA 32 |
| SF8300$^{tet}$ | Tetracycline resistant SF8300 |

**Table S2: List of plasmids used in this study**

| Plasmids used | Notes |
|---|---|
| $pTX_{\Delta}$ | Empty plasmid for constitutive expression[7] |
| $pTX_{\Delta}$ + *gdpP* [SF8300] | Constitutively expressed *gdpP* from SF8300 |
| $pTX_{\Delta}$ + *gdpP* [CRB] | Constitutively expressed *gdpP* from CRB |
| $pTX_{\Delta}$ + *gdpP* [CmTc] | Constitutively expressed *gdpP* from CmTc |
| $pTX_{\Delta}$ + *gdpP* [SRB] | Constitutively expressed *gdpP* from SRB |
| $pTX_{\Delta}$ + *gdpP* (MSSA 4) | Constitutively expressed *gdpP* from MSSA 4 |
| $pTX_{\Delta}$ + *gdpP* (MSSA 7) | Constitutively expressed *gdpP* from MSSA 7 |
| $pTX_{\Delta}$ + *gdpP* (MSSA 9) | Constitutively expressed *gdpP* from MSSA 9 |
| $pTX_{\Delta}$ + *gdpP* (MSSA 13) | Constitutively expressed *gdpP* from MSSA 13 |
| $pTX_{\Delta}$ + *gdpP* (MSSA 19) | Constitutively expressed *gdpP* from MSSA 19 |
| $pTX_{\Delta}$ + *gdpP* (MSSA 21) | Constitutively expressed *gdpP* from MSSA 21 |
| $pTX_{\Delta}$ + *gdpP* (MSSA 22) | Constitutively expressed *gdpP* from MSSA 22 |
| $pTX_{\Delta}$ + *gdpP* (MSSA 24) | Constitutively expressed *gdpP* from MSSA 24 |
| $pTX_{\Delta}$ + *gdpP* (MSSA 25) | Constitutively expressed *gdpP* from MSSA 25 |
| $pTX_{\Delta}$ + *gdpP* (MSSA 27) | Constitutively expressed *gdpP* from MSSA 27 |
| $pTX_{\Delta}$ + *gdpP* (MSSA 28) | Constitutively expressed *gdpP* from MSSA 28 |
| $pTX_{\Delta}$ + *gdpP* (MSSA 32) | Constitutively expressed *gdpP* from MSSA 32 |
| pKOR1 | pKOR1 deletion construct for *gdpP* gene[7] |
| pLL29 | Chromosomal integrative plasmid[8] |

**Table S3: List of primers used in this study**

| Primer | Sequence (5’- 3’) |
|---|---|
| gdpP-P1 | GGGGACAAGTTTGTACAAAAAAGCAGGCTTGTTAATTTTCATTAAAGAGGTTAAAATAATAGCTATAGTTAAAAATATGG |
| gdpP-P2 | TATTCCACCTCTATTCACTTTTTAGAATTATTTTTCATGATTCG |
| gdpP-P3 | AAGTGAATAGAGGTGGAATAATGAAAGTAATTTTTACACAAGATGTTAAAGGTAAAGGTAAAAAAGG |
| gdpP-P4 | GGGGACCACTTTGTACAAGAAAGCTGGGTCTTCAGCTGTTTCATACACTTGTCCTAAGACGTCTCGAATGTCTTTAAAGC |
| gdpP-1 | AAAGGATCCTAAAAAGTGAATAGAGGTGGAATAATG |
| gdpP-2 | ACTAATGACACGTGTTACCATTGAGTTGATTTCC |
| gdpP-3 | ATGGTAACACGTGTCATTAGTCGATGGGCAACTG |
| gdpP-4 | TCGTTCATCACACGTCGTAATGTTGGATCAATG |
| gdpP-5 | AACATTACGACGTGTGATGAACGAAATAGATAAAAAGC |
| gdpP-6 | TAAACGCGTCTTTCATGCATCTTCACTCCTACTTAATTG |
| gdpP -7 | ACGTTTAACGCGACTTGAATCAACAGTGATGTATG |
| gdpP -8 | TGTTGATTCAAGTCGCGTTAAACGTTGTTCTG |

**Table S4: List of the clinical isolates used for measurement of CDA abundance, and their associated *gdpP* mutations.**[9]

| Clinical Strain | *gdpP* associated mutations |
|---|---|
| MSSA 4 | V490E |
| MSSA 7 | F54L |
| MSSA 9 | F54L, P312L |
| MSSA 13 | D105N, P392S, A601E |
| MSSA 19 | A210S |
| MSSA 21 | Q642X |
| MSSA 22 | R504X |
| MSSA 24 | D105N, P392S, V609X |
| MSSA 25 | E486K |
| MSSA 27 | V609D |
| MSSA 28 | T307I |
| MSSA 32 | I203N |

**Table S5: Mutations in *gdpP* and *pbp4* among SRT passaged cultures**

| **Passaged stocks** | **Ceftaroline concentration (mg/L)** | ***gdpP*** | ***Ppbp4+pbp4*** |
|---|---|---|---|
| Day 3 clone 1 | 0.25 | X515 | - |
| Day 3 clone 2 | 0.25 | X402 | - |
| Day 3 clone 3 | 0.25 | X238 | - |
| Day 4 clone 1 | 0.5 | X306 | - |
| Day 4 clone 2 | 0.5 | X306 | - |
| Day 4 clone 3 | 0.5 | X306 | - |

X = missense mutation

**Table S6: MIC (mg/L) of β-lactam drugs**

| Strains | NAF | CTX | FOX | CFZ | CPT |
|---|---|---|---|---|---|
| MW2 | 16 | 64 | 32 | 64 | 0.5 |
| MW2 Δ*gdpP* | 32 | > 256 | 64 | 128 | 1 |
| SF8300 | 16 | 128 | 16 | 16 | 0.5 |
| SF8300$^{tet}$ | 16 | 128 | 16 | 8 | 0.5 |

NAF- nafcillin; CTX- ceftriaxone; FOX- cefoxitin; CFZ- cefazoline; CPT- ceftaroline

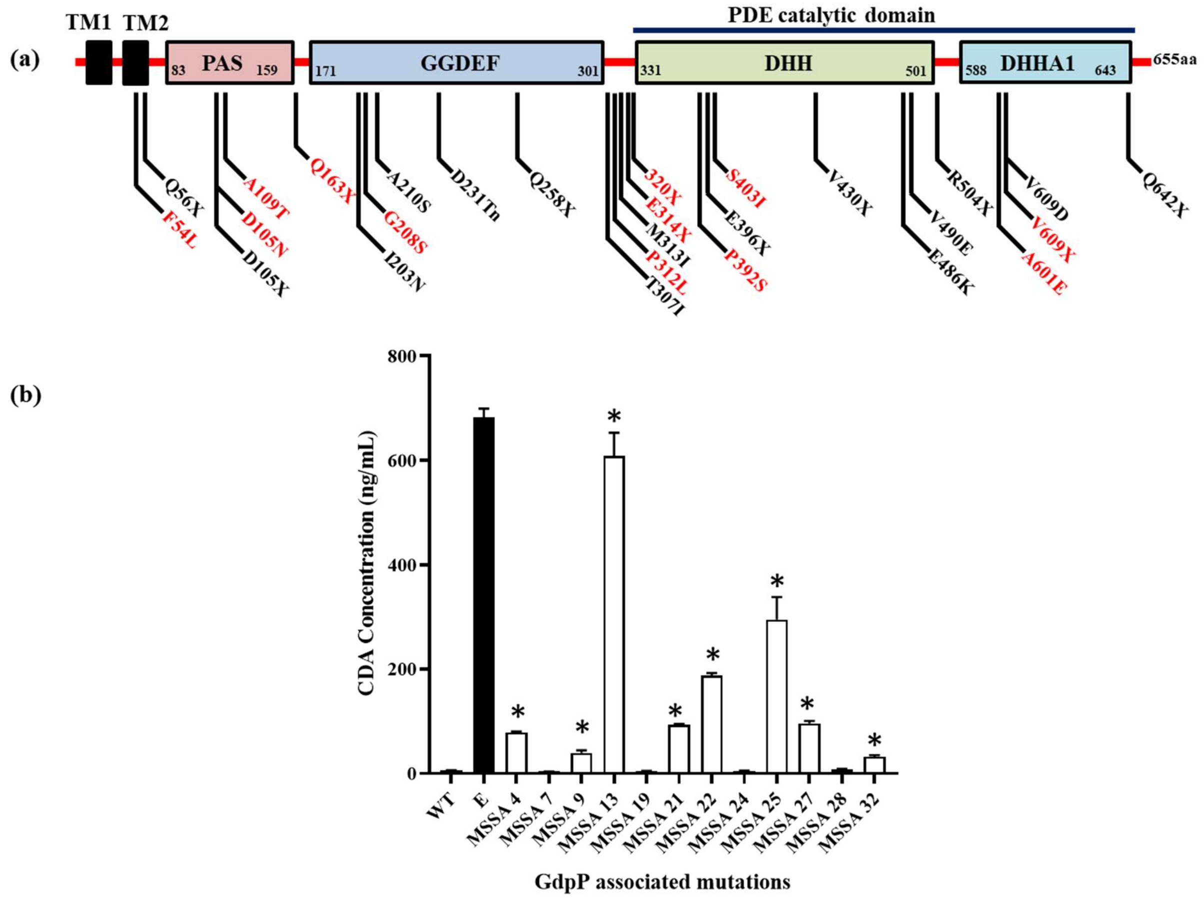


**Figure S1: Clinical strains of S. aureus containing *gdpP* associated mutations.**

**(a)** Schematic diagram showing different domains of GdpP and mutations detected among clinical isolates. Residues in red are present in strains with multiple *gdpP* mutations.

**(b)** Analysis of CDA levels of SF300ex *ΔgdpP* cells containing *gdpP* from clinically isolated strains. The X-axis contains the clinical strains having different mutations associated in the *gdpP* gene. The Y-axis is the CDA concentration. P values: WT versus MSSA 4, WT versus MSSA 13, WT versus MSSA 21, WT versus MSSA 22, WT versus MSSA 27 <0.0001; WT versus MSSA 9, WT versus MSSA 25 = 0.0004; WT versus MSSA 32 = 0.0001.

P values were obtained through two-tailed t-test analysis using GraphPad prism.

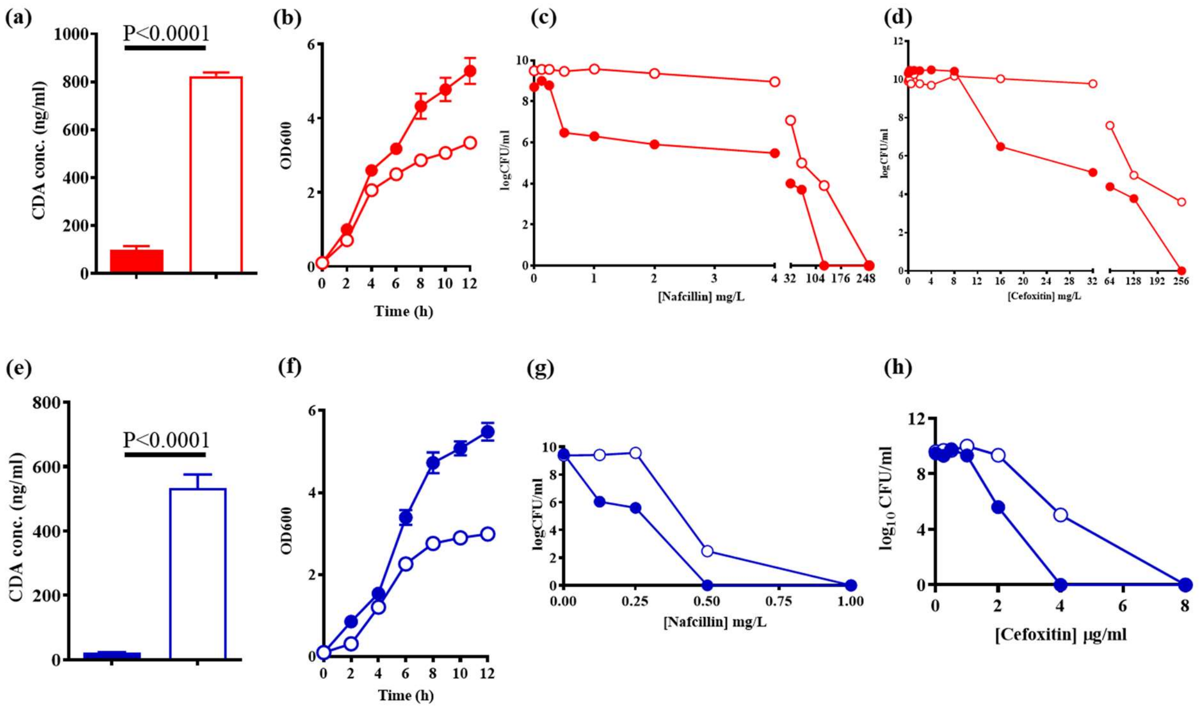


**Figure S2: GdpP phenotypes are not dependent on *mecA*.**

**(a) & (e)** CDA concentrations in the cytosol of SF8300 (closed red) and SF8300ex (closed blue) and their isogenic Δ*gdpP* (open symbols) strains respectively.

**(b) & (f)** Growth curve of SF8300 and SF8300ex wild-type strains (closed red and closed blue circles, respectively) and their isogenic Δ*gdpP* strains (open red and open blue circles, respectively) in TSB media.

**(c) & (g)** Population analysis of SF8300 and SF8300ex wild-type strains (closed red and closed blue circles, respectively) and their isogenic Δ*gdpP* strains (open red and open blue circles, respectively) in nafcillin.

**(d) & (h)** Population analysis of SF8300 and SF8300ex wild-type strains (closed red and closed blue circles, respectively) and their isogenic Δ*gdpP* strains (open red and open blue circles, respectively) in cefoxitin.

P values were obtained through two-tailed t-test analysis using GraphPad prism.

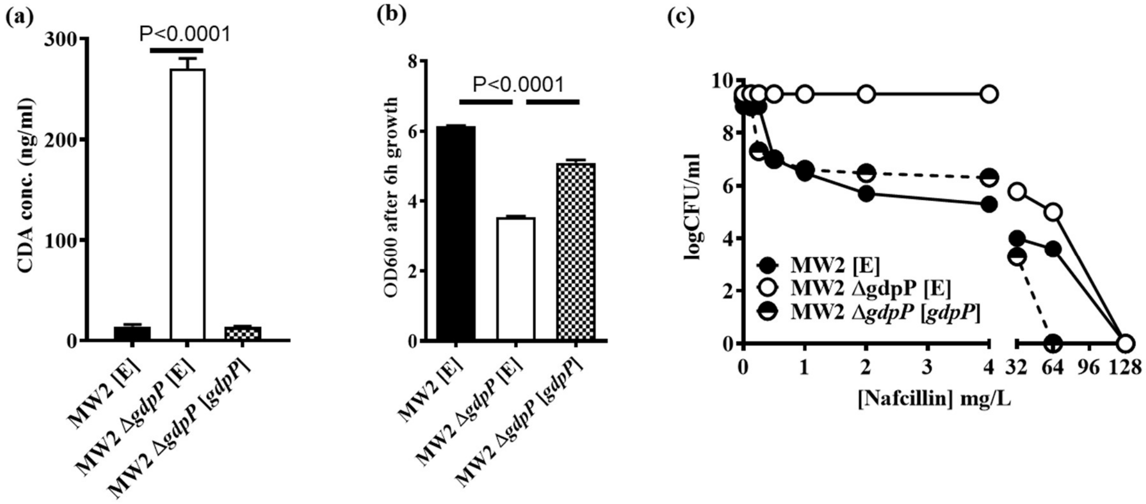


**Figure S3: Complementation with *gdpP* restored wild-type phenotypes in MW2 strains.**

**(a)** CDA levels in the cytosol

**(b)** Growth pattern in TSB media

**(c)** Population analysis with nafcillin

of the complemented MW2 and Δ*gdpP* strains. P values were obtained through two-tailed t-test analysis using GraphPad prism.

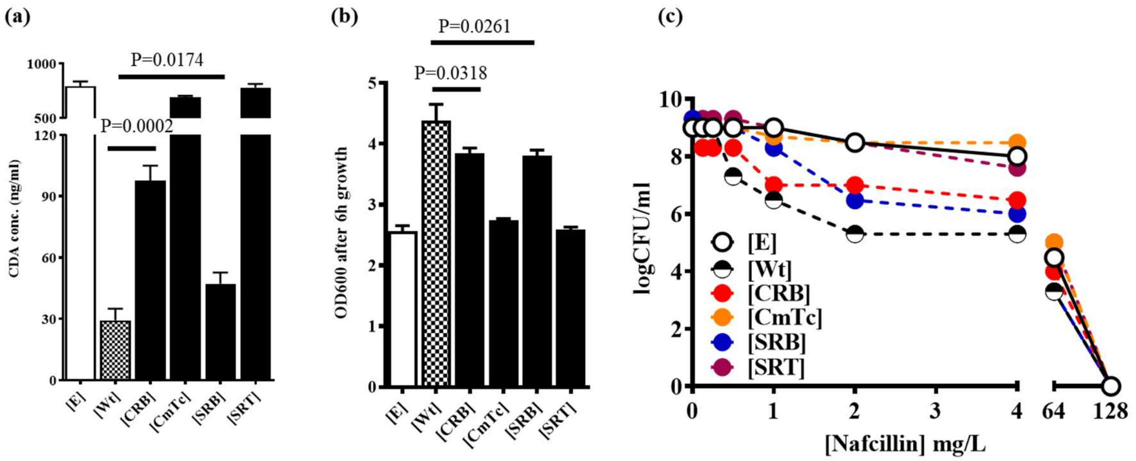


**Figure S4: CDA drives Δ*gdpP* associated phenotypes.**

**(a)** CDA levels in the cytosol

**(b)** OD600 of bacterial cultures in TSB media grown for 6 h

**(c)** Population analysis in nafcillin

of the SF8300 Δ*gdpP* strain complemented with either and empty vector [E], wild-type *gdpP* and mutant *gdpP*s obtained from passaged strains.

P values were obtained through two-tailed t-test analysis using GraphPad prism software and represents analysis between Wt and CRB and Wt and SRB.

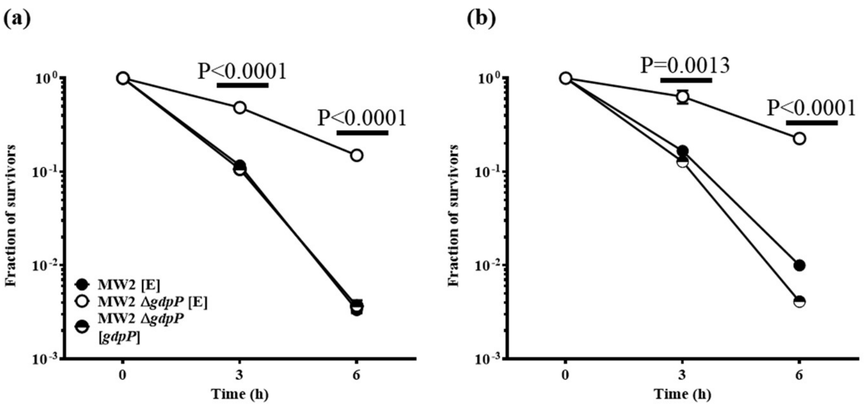


**Figure S5: Deletion of *gdpP* leads to β-lactam tolerance in MW2 strains.**

β-lactam tolerance assay carried out in:

**(a)** nafcillin (128 mg/L)

**(b)** cefoxitin (128 mg/L).

P values represent difference between MW2 [E] and MW2 [*gdpP*] strains and were obtained through two-tailed t-test analysis using GraphPad prism.

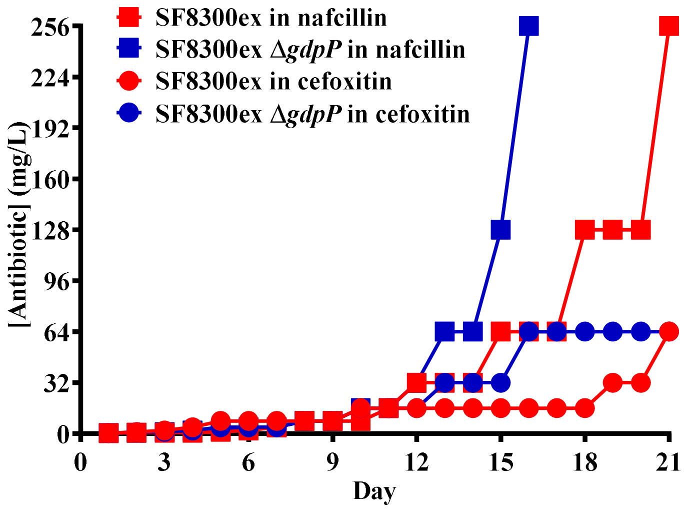


**Figure S6: Passaging of SF8300ex and its Δ*gdpP* strains in nafcillin and cefoxitin.**

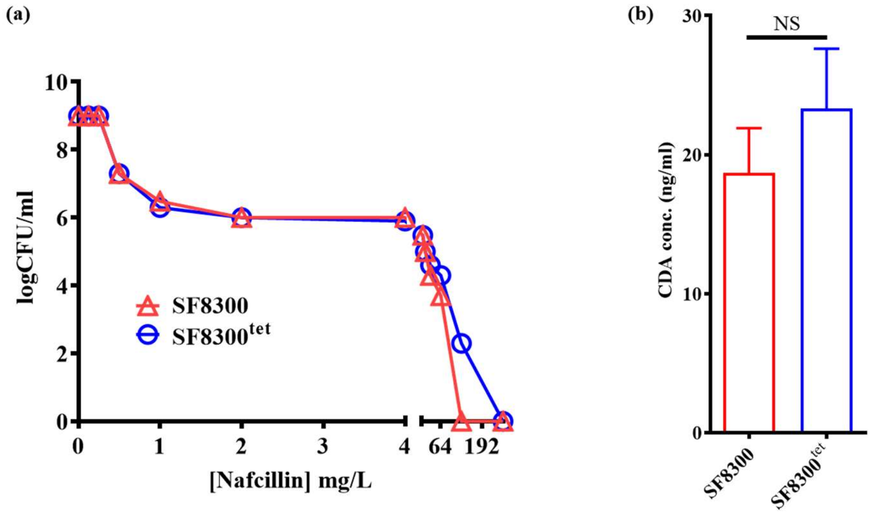


**Figure S7: Loss of *gdpP* function could lead to β-lactam treatment failure.**

**(a)** population analysis with nacfillin

**(b)** CDA concentration in the bacterial cytosol

of SF8300 and pLL29 integrated SF8300$^{tet}$ strains.

P values were obtained through two-tailed t-test analysis using GraphPad prism.

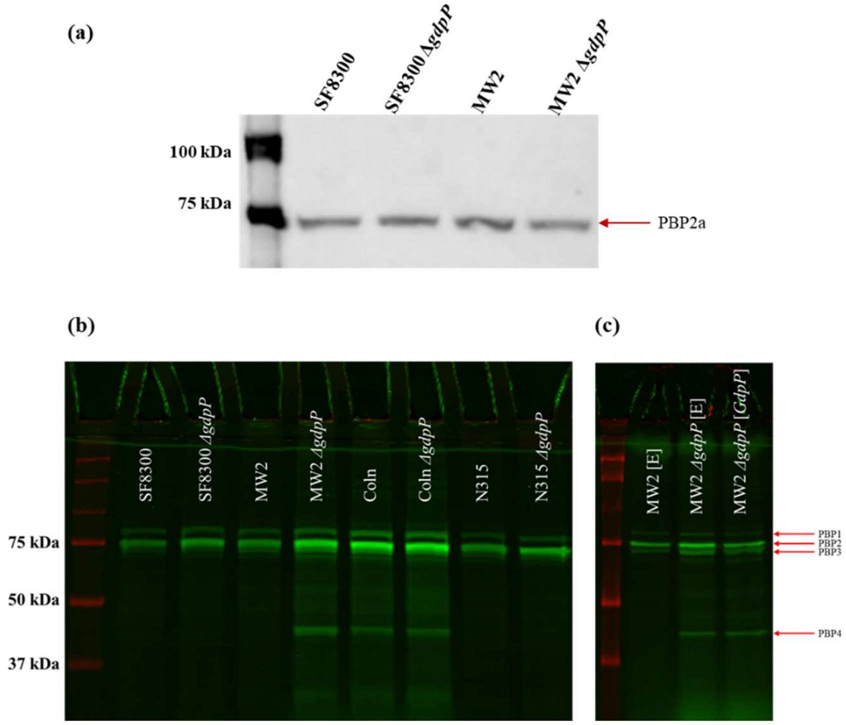


**Figure S8: Deletion of *gdpP* did not have an effect on the expression of PBPs.**

**(a)** Western blots of SF8300, MW2 and their isogenic Δ*gdpP* strains showing PBP2a

**(b)** Bocillin assay showing PBPs in SF8300, MW2, COLn, N315 and their isogenic Δ*gdpP* strain*s*

**(c)** Bocillin assay showing PBPs in MW2, MW2 Δ*gdpP* complemented with an empty vector [E] and MW2 Δ*gdpP* complemented with *gdpP*

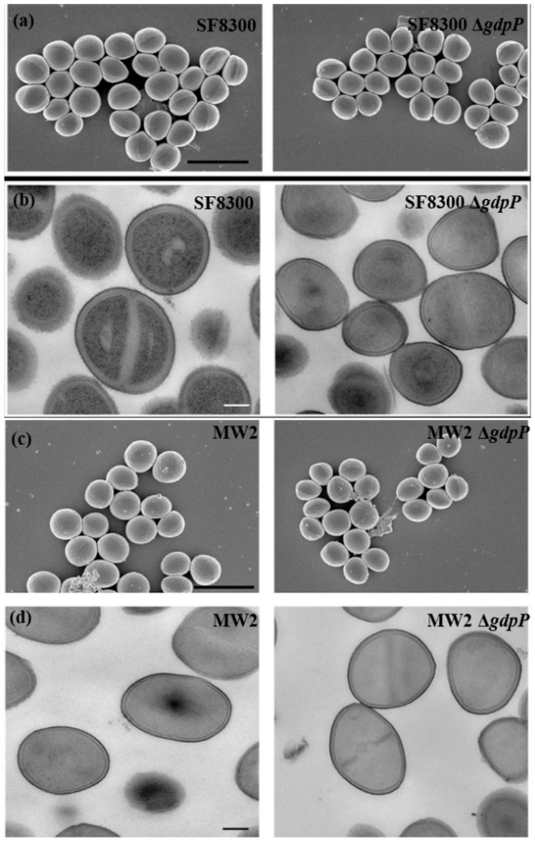


**Figure S9 (a) to (d): Transmission electron microscopic (TEM) and Scanning electron microscopic (SEM) images of SF8300, MW2 and their isogenic Δ*gdpP* strains.**
